# Time-adaptive modulation of evidence evaluation in rat posterior parietal cortex

**DOI:** 10.1101/2025.09.04.674303

**Authors:** Preetham Ganupuru, Adam B. Goldring, Tanner Stevenson, Kendall Stewart, Rishidev Chaudhuri, Timothy D. Hanks

## Abstract

A crucial component of successful decision making is determining the optimal timescale over which to evaluate evidence. For example, when detecting transient changes in the environment, it is best to focus evaluation on the current evidence as opposed to older evidence. However, it is unclear how this adjustment in timescale is achieved in the brain in terms of how the neurons that process evidence adjust their dynamics. To address this question, we used Neuropixel probes to record spiking activity from neurons in the posterior parietal cortex (PPC) of rats performing a free-response auditory change detection task in which subjects evaluate sensory evidence over short timescales to determine when a change occurs in a noisy sensory stream. Consistent with longer timescale temporal integration tasks, we found that PPC neurons modulated their activity by the strength of evidence leading to decisions, were selective for the rats’ choices, and had opposing populations of neurons that were positively versus negatively modulated by evidence. However, in contrast to temporal integration tasks, responses of neurons to individual pulses of evidence were transient, such that the effect of the evidence on activity tapered off over a timescale corresponding to the subject’s behavioral timescale of evidence evaluation. Intriguingly, PPC also exhibited “gain changes” in the influence of evidence as a function of decision time that were consistent with changes in behavioral urgency. In addition, reversible inactivation revealed an important role for PPC in this auditory change detection, such that PPC inactivation altered choice behavior and the timescale over which rats evaluated evidence. Together, our results suggest important contributions of PPC to free-response decisions that involve adjusting timescales of evidence evaluation.

## Introduction

A large body of research has explored how perceptual decision making arises from neural activity across cortical and subcortical regions in tasks that involve the linear integration of evidence over time (Brody and Hanks, 2016; Gold and Shadlen, 2007; Schall, 2019). Among the brain regions studied, pioneering research on the posterior parietal cortex (PPC) has provided considerable insights into generalizable features of neural activity that relate to decision making. In tasks involving multiple alternatives reported through distinct actions, neurons in both primate and rodent PPC steadily increase their firing rates when evidence supports a neuron’s preferred option (Churchland et al., 2008; Erlich et al., 2015; Hanks et al., 2015, 2011; Huk and Shadlen, 2005; Khilkevich et al., 2024; Morcos and Harvey, 2016; Roitman and Shadlen, 2002; Scott et al., 2017; Shadlen and Newsome, 2001; Steinemann et al., 2024). This produces opposing neuron subpopulations whose activity increases for different choice options, with stronger evidence causing a steeper change in activity on average. Further, the effect of individual pieces (or pulses) of evidence on PPC firing rates can be long lasting in a manner that is consistent with linear integration of evidence over time (Hanks et al., 2015; Huk and Shadlen, 2005).

However, there are many situations where linear integration of evidence is not optimal. As two examples, it is not optimal in situations where the environment is unstable or when detecting a transient change in a stimulus. In these situations, it is better to give greater weight to more recent evidence – in other words, to evaluate evidence on shorter timescales. Behavioral results show that subjects can adjust their decision strategy accordingly in these types of situations (Booras et al., 2021; Bronfman et al., 2016; Ganupuru et al., 2019; Glaze et al., 2015; Gold and Stocker, 2017; Harun et al., 2020; Levi et al., 2018; Ossmy et al., 2013; Piet et al., 2018; Radillo et al., 2017; Tsetsos et al., 2012). In the domain of perceptual decision making, it has been shown that human subjects performing a change detection task can flexibly adjust the timescale of evaluation of past evidence depending on the type of decision they make (Ganupuru et al., 2019) and that these changes are adapted to task demands (Booras et al., 2021; Harun et al., 2020). The first-order question we seek to address in this study is how PPC dynamics differ when rats perform a similar change detection task that involves relatively short timescales of evidence evaluation.

Another key factor influencing decision dynamics over time is urgency—a time-dependent drive to commit to a choice. Behavioral studies suggest urgency can potentially modulate response timing and accuracy (Cisek et al., 2009; Drugowitsch et al., 2012; Hanks et al., 2014; Kira et al., 2025; Murphy et al., 2016; Reddi and Carpenter, 2000; Shinn et al., 2020; Thura and Cisek, 2016). It has been further suggested that PPC neural responses reflect urgency in the form of evidence-independent neural activity that increases as time elapses during decision formation (Churchland et al., 2008; Hanks et al., 2014; Purcell and Kiani, 2016). Another not mutually exclusive possibility is that decision urgency could result from gain changes on how evidence impacts decision circuits such that the influence of equivalent evidence increases as a function of decision time. Using the pulsatile delivery of evidence in the present work, we seek to directly test the idea that there may be gain changes for the impact of evidence as a function of decision time.

Finally, there is considerable debate about the causal role of PPC in perceptual decision making. In studies involving linear integration of evidence over time, microstimulation of primate PPC can cause small but reliable effects on decision making (Fetsch et al., 2014; Hanks et al., 2006). However, in similar paradigms, there have been mixed results with pharmacological inactivation and optogenetic perturbation of primate and rodent PPC, with some experiments finding effects and others not (Akrami et al., 2018; Erlich et al., 2015; Jeurissen et al., 2022; Katz et al., 2016; Licata et al., 2017; Yao et al., 2020; Zhou and Freedman, 2019). A possible reconciling perspective comes from the observation that effects have been found most consistently in free response tasks rather than cued response tasks. The change detection task we use here involves a free response without lengthy integration of evidence, thereby allowing us to test whether inactivation of PPC affects decision making in this situation.

## Results

### Behavioral performance

We trained rats to perform a free-response auditory change detection task that was adapted from previous studies involving human subjects (Johnson et al., 2017). Rats are cued with a light to insert their noses into a nose port of an operant apparatus, which initiates a stream of broadband auditory pulses (“clicks”) generated via a Poisson process. Starting at a baseline rate of 20 Hz, the generative click rate may increase at a random time by a variable magnitude, to which subjects must respond within 0.8 s by withdrawing their nose from the port. A timely withdrawal in response to a change is a *hit*, which leads to a water reward delivered from a separate port on the side of the apparatus. Failure to withdraw in time results in a *miss*, with no reward. Withdrawal in the absence of a change is a *false alarm*, also denying a reward. Finally, on a subset of trials, no change in stimulus rate occurs; during these “catch” trials, the rat must maintain port fixation until the stimulus ends, resulting in a *correct rejection* accompanied by a water reward (**Figure 1A**).

**Figure 1:**
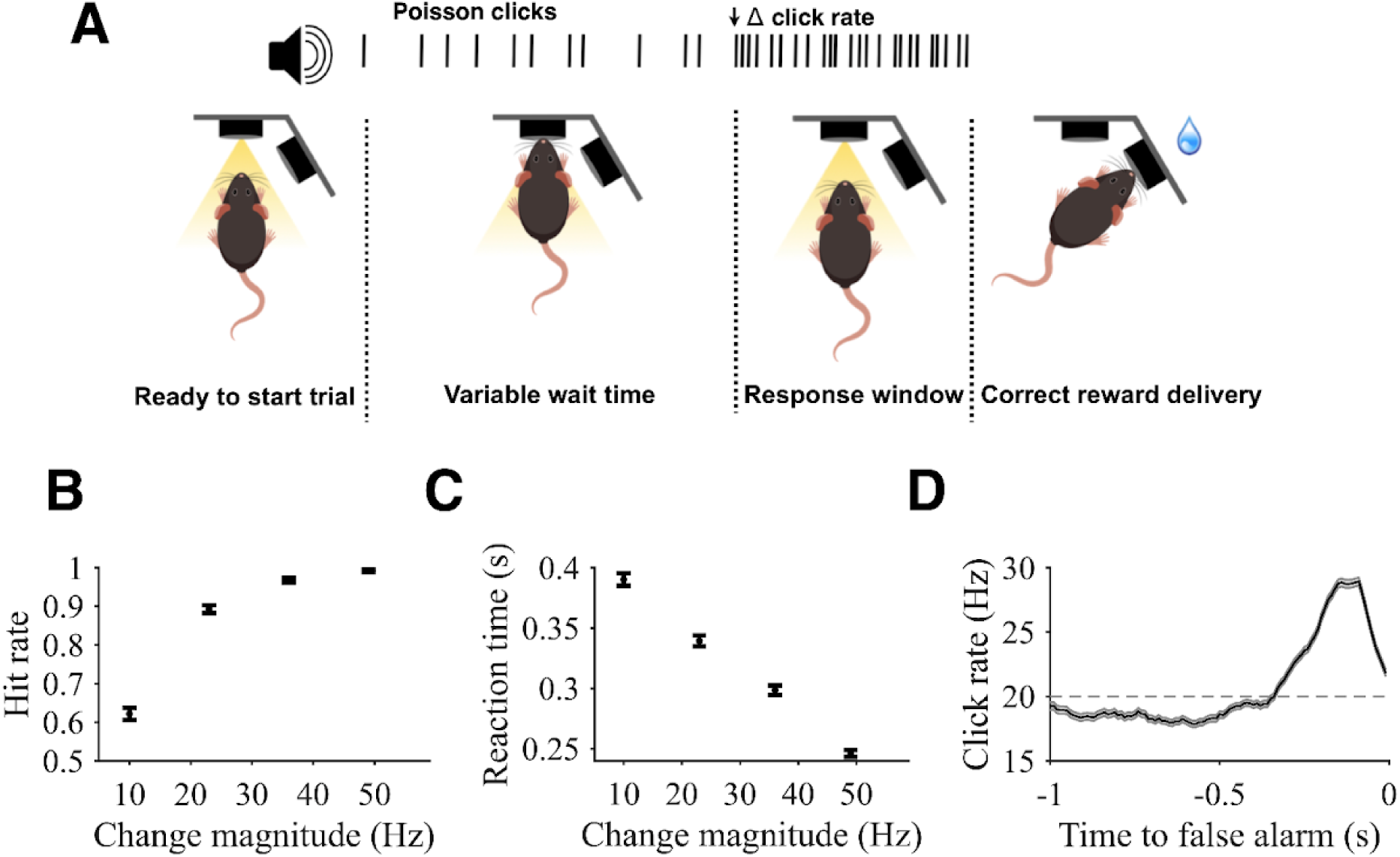
Task design and subject performance. **A)** A trial begins when a rat pokes an illuminated port in the apparatus with its nose. Port engagement triggers initiation of a stream of Poisson-generated auditory pulses (“clicks”) at a baseline generative rate of 20 Hz. At a random point (marked by black arrow in this example) in 70% of trials, the generative rate increases by a variable magnitude of 10 Hz, 23 Hz, 36 Hz, 49 Hz. After the rate change, rats have 0.8 s to withdraw their nose from the initial port in order to receive a water reward from a second port. In the other 30% of trials, no change occurs, and rats must maintain port fixation for the full duration of the stimulus to receive a water reward. **B)** Combined hit rates as a function of change magnitude. Hit rate was calculated excluding false alarm trials (i.e. only including trials in which the rat was presented with a change). **C)** Reaction times for hits as the time of center port withdrawal following a change. **D)** Psychophysical reverse correlation (PRC). Calculated as the average local click rate preceding false alarms. Horizontal line shows the 20 Hz baseline rate. For all plots, error bars and error shading show SE. N = 4 rats.

Like humans, rats consistently detected changes, achieving higher hit rates with higher change magnitudes (Regression coefficient: 0.11 +/- 0.006, p < 0.001; **Figure 1B**). Higher change magnitudes also yielded faster reaction times (Regression coefficient: -3.65 +/- 0.21, p < 0.001; **Figure 1C**). These metrics suggest rats perform the task by evaluating the presented sensory evidence, and greater quantities of evidence lead to faster and more accurate choices. In addition, while rats were prone to false alarms (average false alarm rate was 0.45 +/- 0.006), they were also able to withhold responses for the entire duration of catch trials (average correct rejection rate of 0.39 +/- 0.01).

To better gauge the strategy rats employ on this task, we conducted psychophysical reverse correlation (PRC) analyses using false alarm trials to assess how rats evaluated evidence over time prior to their decision to withdraw their nose from the port. PRC traces were constructed by convolving click times with causal half-Gaussian filters (sigma = 0.05 s) and aligning the result to the time of the false alarm response. Because the Poisson stimulus varies around a generative mean from moment to moment, this analysis reveals how false alarms are related to stochastic fluctuations in click rate as a function of time. Therefore, the PRC method provides insight into the timescale and strength of evidence contributing to decision formation (Ganupuru et al., 2019; Okazawa et al., 2018).

As expected, the false alarm PRC exhibited an upward fluctuation in click rate prior to false alarms, indicating false alarms were generally associated with brief increases in click rate (**Figure 1D**). The period of this upward fluctuation extended 0.36 +/- 0.02 s prior to the rat’s choice, consistent with a short timescale of evidence evaluation. The mean increase of the click rate from 50 to 350 ms before the false alarm was 5.8 +/- 0.2 Hz (p < 0.001). The PRC thus suggests a behavioral strategy on the part of the rats of evaluating evidence over relatively brief intervals while discounting older evidence to detect increases in stimulus rate, consistent with task demands.

### Evidence-modulated neural responses in PPC

After characterizing the rats’ dynamics of evidence evaluation at the behavioral level, we sought to examine their neural underpinnings. The spiking responses of 1077 single units in PPC contralateral to the reward port were recorded using Neuropixels probes while rats performed the change detection task, and those neurons were then assessed for task-related activity (**Figure 2**). We found 258 neurons (24%) with significant modulation during decision formation as assessed by comparison of firing rates during the change period versus the baseline period (from stimulus onset until the change). Of those 258 neurons, 92 were positively modulated and 166 were negatively modulated leading up to the decision. We focus first on the neurons that were positively modulated (**Figure 3**).

**Figure 2:**
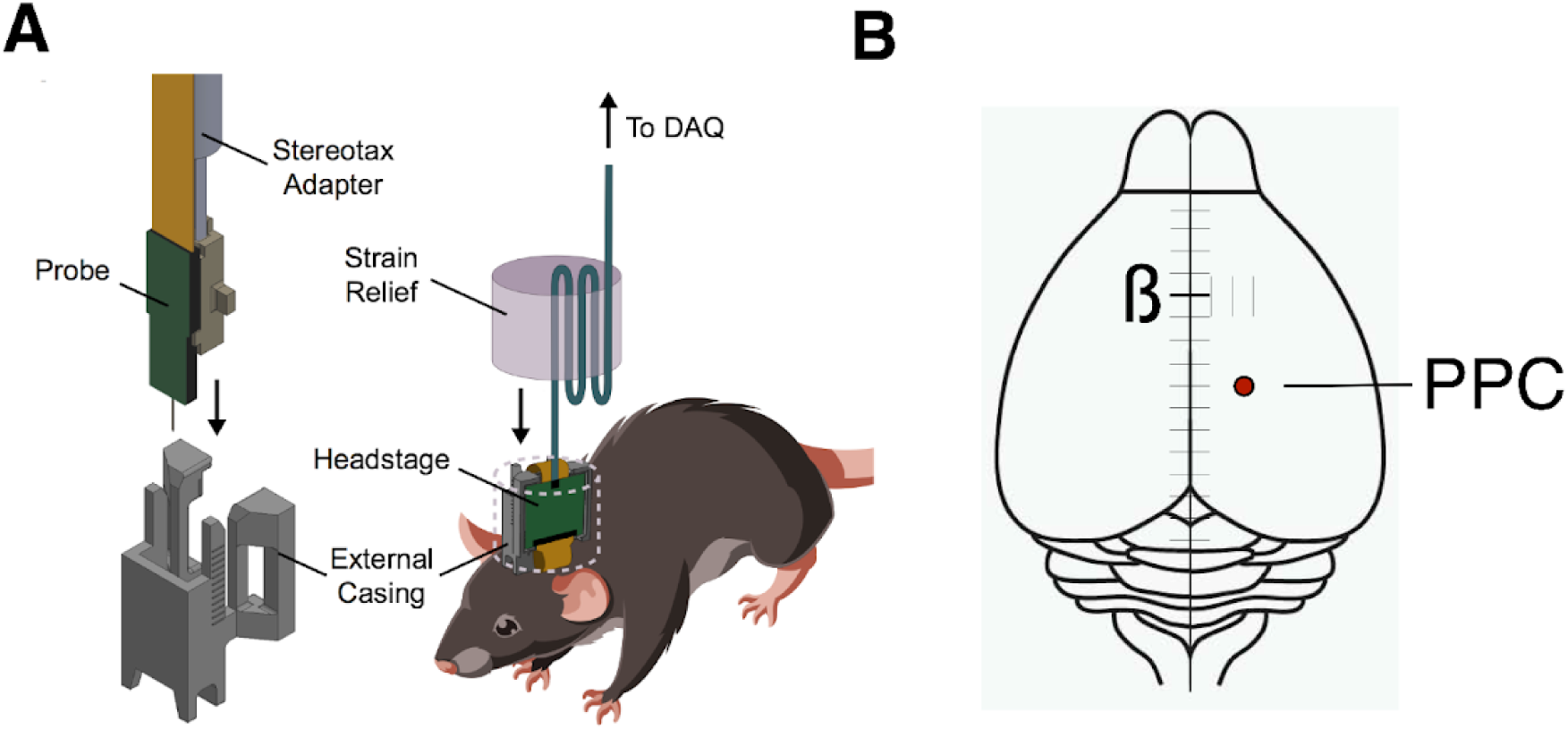
Recording methods. **A)** For each rat, a Neuropixels recording probe was implanted using a custom designed assembly that allowed use of a strain-relieving attachment for the tether. **B)** Probes were targeted stereotaxically to PPC (-3.8 to -3.95 mm AP, 2.2 mm ML).

**Figure 3:**
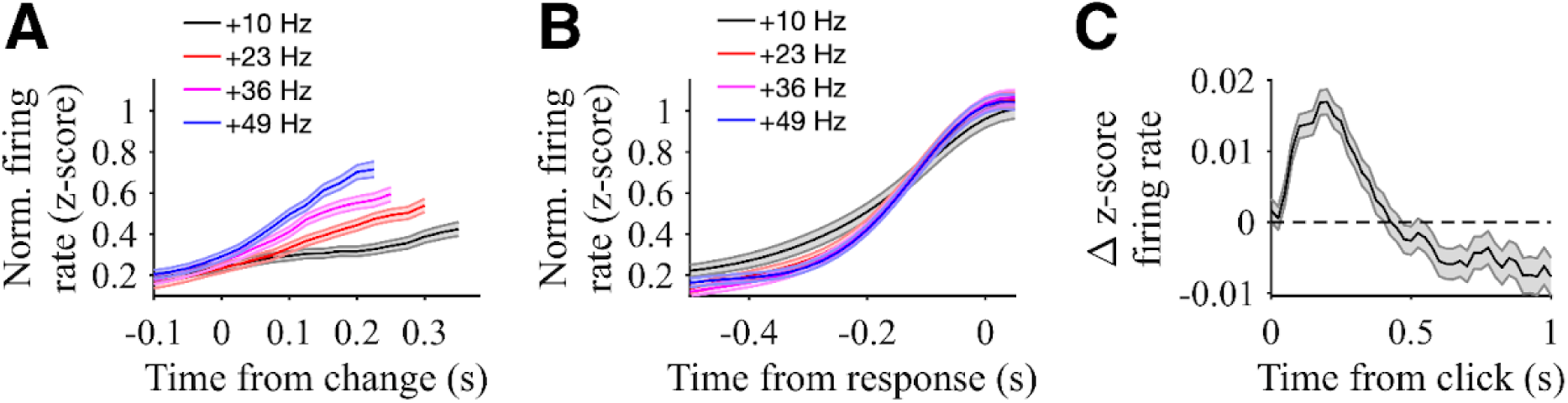
Responses of PPC neurons positively modulated by evidence. **A)** Population responses of neurons with significant positive modulation by evidence strength aligned to time of stimulus change (same rats as in Figure 1). Traces are normalized by the pre-change firing rates (from stimulus onset to change time) and sorted by change magnitude. Responses are plotted to the median reaction time for each change magnitude. Shading shows SE. N = 92 neurons. **B)** Population responses for hit trials of the same neurons aligned to the time of hit response. Traces are normalized by the pre-change firing rates and sorted by change magnitude. Shading shows SE. N = 92 neurons. **C)** Population click-triggered response of the same neurons. Residual z-scored responses aligned to each click are averaged across clicks for all neurons. Only clicks during the pre-change period are included, and neural responses are masked after the change (for hit and miss trials), movement response (for false alarm trials), or end of stimulus (for correct rejection trials). Shading shows SE. N = 92 neurons.

The average activity of these neurons was modulated by the strength of sensory evidence when aligned to the stimulus change (**Figure 3A**). Activity ramped up for all stimulus conditions with greater click rate increases associated with larger average firing rates after the stimulus change when aligned to that change (Regression coefficient: 0.0059 +/- 0.0008, p<0.001). Importantly, as described above, subjects had faster reaction times for trials with greater click rate increases, which can contribute to the differences in firing rate aligned to the stimulus change. Thus, it is valuable to also consider how neural activity compared across stimulus conditions aligned to the time of the decision response. Differences between stimulus conditions were less evident when aligned to the decision response (**Figure 3B**). A common level of activity was reached leading up to the decision response, and there was not a significant modulation when considering the time period from 150 ms to 50 ms before the decision response (Regression coefficient: 0.0001 +/- 0.0011, p=0.91). Taken together, these results suggest that, in this free-response change detection task, the activity of positively modulated PPC neurons increases at a rate proportional to the amount of evidence presented over time, until reaching a common level of activity preceding the response. This trend suggests that this subpopulation of PPC neurons may play a role in integrating evidence until a decision commitment threshold is hit; however, it does not give insight into the underlying timescale of that integration.

To directly test the hypothesis that PPC neural dynamics reflect shorter timescales of evidence integration used in our change detection, we computed a click-triggered average of the change in neural activity for positively modulated neurons to single pulses of evidence – that is, to single clicks (**Figure 3C**). The resulting average showed a transient increase in firing rate following clicks (average increase from 50 ms to 350 ms after a click was 0.0069 +/- 0.0015 z-score, p < 0.01). This analysis is applied only to clicks during the baseline pre-change period, thus excluding the post-change intervals analyzed above and providing separate verification of the influence of clicks on neural activity. Notably, the duration of this increase extended less than 500 ms following the click, which is broadly consistent with the short timescale of evidence integration for this task. This timescale also differs from that reported in similar analyses conducted on positively modulated PPC neurons in an auditory Poisson discrimination task that involved longer timescales of evidence integration (Hanks et al., 2015), suggesting that PPC dynamics are flexible based on task demands. Intriguingly, the click-triggered average firing rate change in our task also fell below baseline after the transient increase, suggesting that clicks that do not lead up to an immediate decision eventually have a negative influence on the activity of positively modulated neurons. We consider this finding further in the discussion section below.

### Gain changes of PPC neural responses across decision time

Because comparison of our results to previous work suggests flexibility of PPC dynamics for different task demands, we next sought to answer whether PPC responses are adaptive across time within a single task, again focusing on positively modulated neurons. In particular, we wanted to test whether there were changes in the influence of sensory evidence on PPC activity that aligned with modulations in how stimuli influenced decision-making behavior. We first measured whether the time of a false alarm response changes the shape of the psychophysical reverse correlation (PRC). We split false alarm trials into whether the nose withdrawal was ‘early’ (<1.5 s after stimulus start) or ‘late’ (>1.5 s after stimulus start) and calculated separate PRCs for each group (**Figure 4A**). Trials ending with an earlier false alarm were associated with a significantly larger PRC increase compared to trials ending with a later false alarm (mean difference of 1.88 +/- 0.18 Hz from 350 ms to 50 ms before the false alarm, p < 0.001). This suggests that “more evidence” is required to trigger an early false alarm compared to a later false alarm, and is consistent with increasing decision urgency and/or decreasing decision bounds as a function of elapsed decision time.

**Figure 4:**
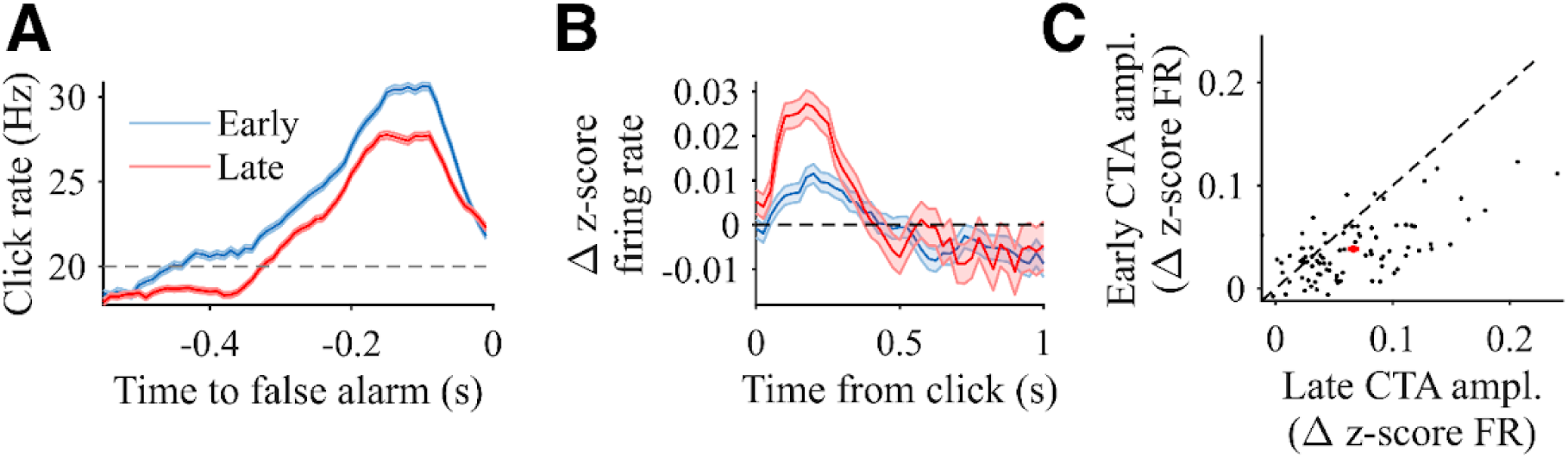
Behavioral and PPC neural responses for early and late stimulus periods. **A)** Psychophysical reverse correlation (PRC) for the same rats as Figure 1, calculated as the average local click rate preceding false alarms and sorted by time of the false alarm. The blue trace corresponds to trials in which a false alarm occurred within 1.5 s of the stimulus start, and the red trace corresponds to trials in which a false alarm occurred later than 1.5 s after stimulus start. Horizontal line shows the 20 Hz baseline rate. Error shading shows SE. N = 3992 trials. **B)** Population click-triggered responses of the positively modulated neurons, sorted by time of the click. The blue trace corresponds to clicks presented within 1.0 s after stimulus start, and the red trace corresponds to clicks presented at least 1.0 s after stimulus start. For both, only clicks during the pre-change period are included, and all neural responses are masked after the change (for hit and miss trials), movement response (for false alarm trials), or end of stimulus (for correct rejection trials). Shading shows SE. N = 92 neurons. **C)** Click-triggered average (CTA) magnitude for each neuron comparing early clicks (y-axis) versus late clicks (x-axis), defined in the same manner as panel B. Each black point corresponds to one individual neuron. N = 92 neurons. The red point shows the average across neurons with error bars corresponding to the SE.

We next tested whether there was any corresponding difference in the influence of evidence on PPC activity as a function of elapsed trial time. To assess this, we returned to the click-triggered analysis but here applied separately to clicks that occur earlier versus later in a trial (**Figure 4B**). Considering the approximately 500 ms average of positive influence of a click on PPC activity, we used a split point that was 500 ms earlier for this analysis as that used for the behavioral comparison in Figure 4A, splitting ‘early’ and ‘late’ clicks (those <1 s versus >1 s from stimulus start, respectively).

Intriguingly, this analysis reveals a change in the magnitude of the click-triggered transient increase, with a significantly smaller influence of earlier compared to later clicks (average difference from 50 ms to 350 ms after a click was 0.0065 +/- 0.0015 z-score, p<0.05). Similar results are apparent when comparing the click-triggered influence of earlier versus later clicks for individual neurons (**Figure 4C**). These analyses suggest a “gain change” in PPC as a function of elapsed decision time, such that early evidence has lesser influence on PPC activity than later evidence. A gain change of sensory evidence of this nature could provide a mechanism by which more evidence is required for an earlier decision compared to a later decision, as we have shown at a behavioral level. In summary, adjustments in the influence of evidence at the behavioral level as a function of elapsed decision time align with gain changes of PPC neurons during evidence evaluation.

### Negatively modulated neural responses in PPC

As described above, in addition to neurons in PPC that were positively modulated during the decision task, an even larger number were negatively modulated. Not surprisingly, the population activity of these neurons was modulated by the strength of sensory evidence when aligned to the stimulus change in an opposing manner to positively-modulated neurons (**Figure 5A**). Activity ramped down for all stimulus conditions with greater click rate increases associated with larger decreases in neural activity after the stimulus change (Regression coefficient: -0.0015 +/- 0.0005, p<0.001). Unlike the positively-modulated neurons, there were differences in negatively modulated activity leading up to the decision response (**Figure 5B**), with significant modulation as a function of stimulus strength in the time period from 150 ms to 50 ms before the decision response (Regression coefficient: 0.0015 +/- 0.0005, p<0.001).

**Figure 5:**
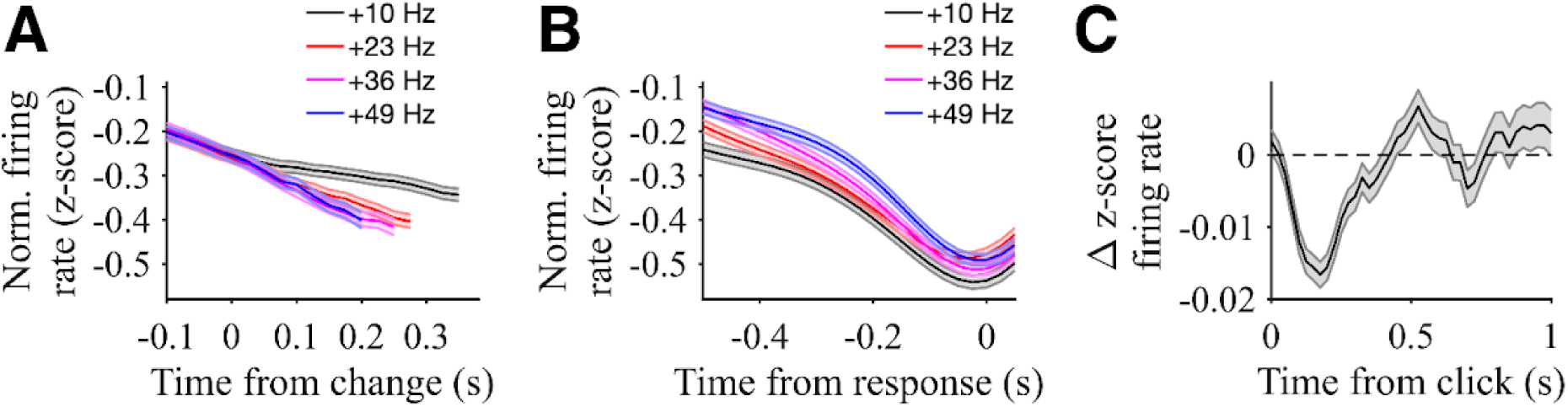
Responses of PPC neurons negatively modulated by evidence. **A)** Population responses of neurons with significant negative modulation by evidence strength aligned to time of stimulus change (same rats as in Figure 1). Traces are normalized by the pre-change firing rates and sorted by change magnitude. Responses are plotted to the median reaction time for each change magnitude. Shading shows SE. N = 166 neurons. **B)** Population responses of the same neurons aligned to the time of hit response. Traces are normalized by the pre-change responses and sorted by change magnitude. Shading shows SE. N = 166 neurons. **C)** Population click-triggered response of the same neurons focusing on “late clicks” at least 1.0 second after stimulus start. Residual normalized responses aligned to each click are averaged across clicks for all neurons. Only clicks during the pre-change period are included, and neural responses are masked after the change (for hit and miss trials), movement response (for false alarm trials), or end of stimulus (for correct rejection trials). Shading shows SE. N = 166 neurons.

In our change detection task, subjects need to determine at each moment in time whether to 1) report a change or 2) not (yet) report a change. Thus, one possibility is that negatively modulated neurons are associated with suppression of decision responses and are therefore inhibited by evidence that supports a decision response. Therefore, we also performed click-triggered analyses for this population of neurons (**Figure 5C**), focusing on clicks occurring at least 1 s after the stimulus, which had the most robust influence on positively modulated neurons. Consistent with other results, the negatively modulated neurons exhibited a transient decrease in activity following clicks (average decrease from 50 ms to 350 ms after a click was -0.0050 +/- 0.0015 z-score, p<0.01). Taken together, these results show a neural dichotomy with opposing subpopulations within PPC in our “go/no go” change detection task. This shares many characteristics with PPC neural responses that have been differentiated on a lateral dichotomy in past decision making studies that employed tasks requiring subjects to make a lateralized choice involving some form of leftward or rightward movement (Churchland et al., 2008; Erlich et al., 2015; Hanks et al., 2015, 2011; Huk and Shadlen, 2005; Roitman and Shadlen, 2002; Shadlen and Newsome, 2001; Steinemann et al., 2024).

### Causal role of PPC in evidence evaluation in change detection

Finally, we sought to determine how disruptions of PPC impacted change detection decisions. Although examinations of PPC neural dynamics have prominently posited a role in evidence integration for decision making, some prominent studies involving perturbation of PPC have suggested a lack of contribution (Erlich et al., 2015; Katz et al., 2016). However, these studies commonly featured a decision time controlled by the environment rather than by the subject. Indeed, PPC has been reported as having a causal role in situations where the animal–and not the environment–controls the time of decision (Hanks et al., 2006; Zhou and Freedman, 2019), as in the case of our task. Therefore, it is plausible that, even though PPC may not be necessary in the accumulation of sensory evidence for perceptual decisions over long timescales, it may play a role in decision processes in situations where the animal decides how and when to respond to incoming stimuli.

To test this hypothesis, we reversibly inactivated the PPC in the change detection task via infusions of the GABA agonist muscimol. Then, we compared task performance between muscimol sessions and sessions in which subjects were infused with vehicle saline solution. With muscimol inactivation of PPC, rat subjects had lower hit rates on average compared to control sessions (Regression coefficient for logistic shift of hit rate function: β = -0.68 +/- 0.14, p<0.001; **Figure 6A**). They also had fewer false alarms (Vehicle: 0.42 +/- 0.01, Muscimol: 0.31 +/- 0.01, p<0.001; **Figure 6B**), suggesting a general reduction in propensity to commit to a report or sluggishness in decision commitment. Consistent with that, reaction times on hit trials also increased with PPC inactivation compared to control sessions (Regression coefficient for shift of RT function: 24.1 +/- 6.6 ms, p<0.001; **Figure 6C**).

**Figure 6:**
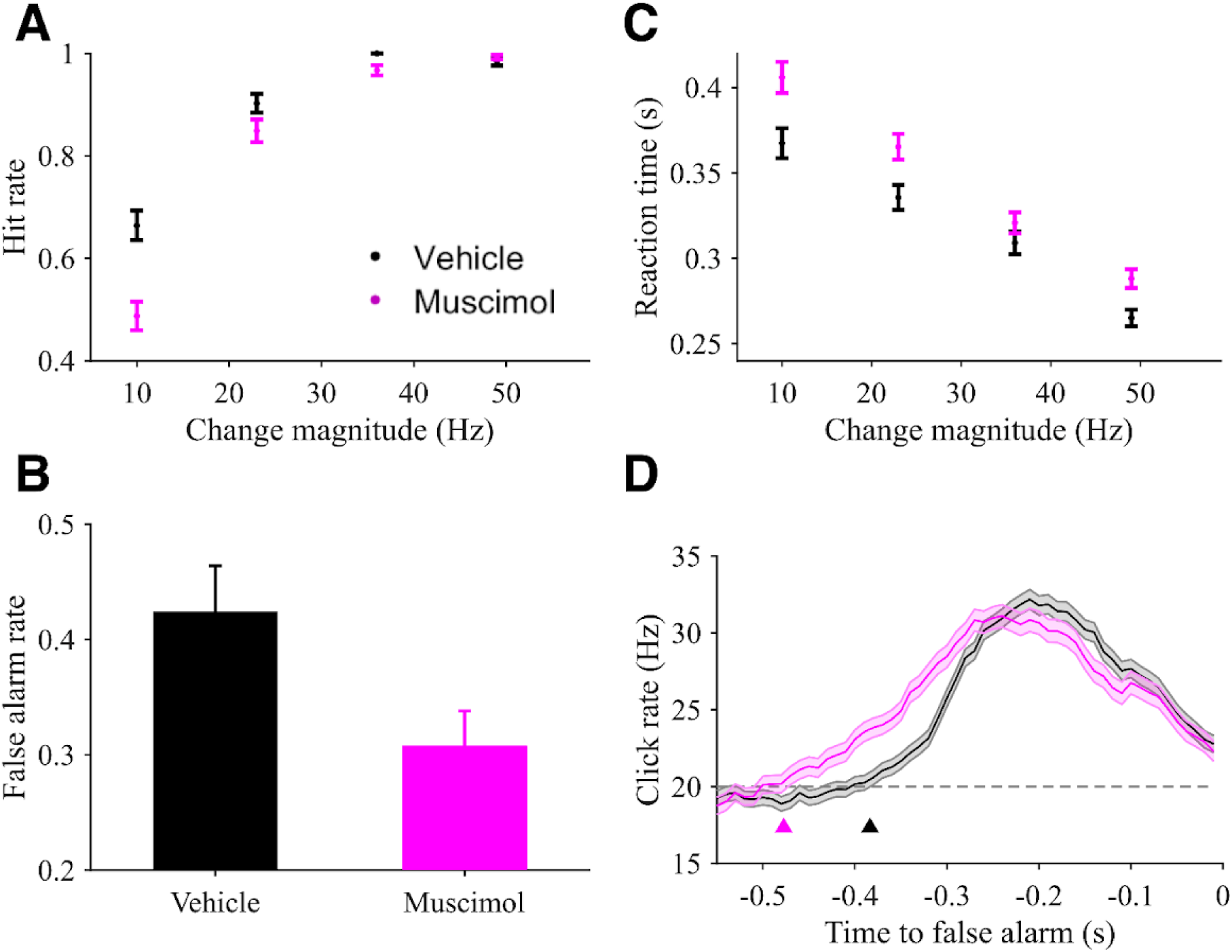
Rat behavioral performance following muscimol inactivation of PPC. **A)** Hit rates as a function of change magnitudes comparing sessions with PPC infusion of vehicle (black) versus muscimol (magenta). Error bars show SE. **B)** False alarm rates during task performance following PPC infusion of vehicle (black) and muscimol (magenta). Error bars show SE. **C)** Correct trial reaction times for vehicle (black) and muscimol (magenta) sessions. **D)** Psychophysical reverse correlation (PRC) for vehicle (black) and muscimol (magenta) sessions. The horizontal line shows the 20 Hz baseline rate. Arrowheads show estimated start time of the increase in the PRC. Error shading shows SE. N = 3 rats for all panels.

These behavioral changes may be explained by multiple alternative mechanisms, including changes in decision bound and/or changes in dynamics of the decision process. To shed further light on this, we again used psychophysical reverse correlation (PRC) analyses as described above. In contrast to the decrease in PRC peak magnitude with elapsed trial time, the PRC for muscimol sessions had a leftward temporal shift when compared to that of control sessions, resulting in an earlier start of the average click rate increase (Muscimol: -0.48 +/- 0.02 s, Vehicle: -0.38 +/- 0.01 s, p<0.001; **Figure 6D**). This difference in PRC suggests a more sluggish timescale of evidence evaluation when PPC is inactivated, consistent with the influence on choices and reaction times.

## Discussion

Our results demonstrate that the rat posterior parietal cortex (PPC) dynamically modulates evidence evaluation in a time-adaptive manner to suit the behavioral demands of a free-response change detection task. In contrast to classic temporal integration tasks in which evidence is accumulated over seconds (Brody and Hanks, 2016; Gold and Shadlen, 2007; Schall, 2019), we observed more transient, short-lived neural responses to individual pulses of sensory evidence in PPC for the change detection task that involved correspondingly brief behavioral evidence evaluation timescales. This shows that PPC circuits are capable of dynamically shaping their evidence evaluation window based on task demands.

The influence of evidence on PPC activity also varied systematically with elapsed decision time. Later-occurring evidence pulses elicited stronger neural responses than early ones, consistent with a time-dependent gain modulation. These gain changes paralleled behavioral findings that less evidence was required to elicit late decisions than early ones—a hallmark of increasing decision urgency (Cisek et al., 2009; Drugowitsch et al., 2012; Hanks et al., 2014; Kira et al., 2025; Murphy et al., 2016; Reddi and Carpenter, 2000; Shinn et al., 2020; Thura and Cisek, 2016). These observations suggest that urgency signals may be produced, at least in part, through changes in the scaling or gating of evidence (Standage et al., 2011).

Even though the change detection task involved only a single active response option, some PPC neurons were positively modulated by evidence while other PPC neurons were negatively modulated by evidence. This suggests a broader principle of functional opponency (Churchland and Ditterich, 2012), even in the absence of lateralized left-right choice paradigms. In the context of the go/no-go structure of our task, these populations may instead support response initiation and suppression, respectively. Such a scheme could allow the brain to explicitly balance the urgency to respond against the risk of premature action, with positively modulated neurons driving “go” signals and negatively modulated neurons involved in a dynamic threshold or inhibitory gate on decision commitment, reminiscent of omnipause neurons in saccadic control (Cohen and Henn, 1972; Evinger et al., 1982; Keller, 1974; Luschei and Fuchs, 1972).

Our results also clarify the conditions under which PPC exerts a causal influence on decision making. Inactivation of PPC via muscimol significantly altered performance, slowing responses and resulting in fewer “detect” (hit and false alarm) reports. Inactivation also shifted the temporal profile of the psychophysical kernel, suggesting sluggish evaluation of evidence. These changes are consistent with a role for PPC in modulating decision timing and dynamics. It is also noteworthy that these effects emerge in a free-response task in which animals determine when to respond. This aligns with prior evidence that PPC’s contributions are most apparent when decisions require internal control of response timing (Hanks et al., 2006; Zhou and Freedman, 2019) rather than in cue-driven paradigms where response timing is externally imposed (Erlich et al., 2015; Katz et al., 2016).

In summary, our findings reveal a dynamic and time-adaptive role for PPC in perceptual decision making. Rather than having rigid dynamics, PPC flexibly shapes the timescale and gain of evidence evaluation to support task demands.

## Methods

### Subjects

A total of 7 male Long-Evans rats aged 1-2 years were used for this study. 4 rats were used for neural recordings and 3 rats were used for the inactivation experiment. Rats were on a water schedule outside of behavioral sessions such that 30 minutes after each behavioral session, they were given free access to water for one hour.

### Apparatus

Tasks were programmed and run in MATLAB (Mathworks) using Bpod (Sanworks) for real-time control and measurement of behavioral output. Operant chambers used for behavioral data collection consisted of three ports. Each port contained an infrared LED beam that detects rat nose insertion upon obstruction of the beam, as well as an LED light that could be used as a cue for the rats. In addition, speakers were mounted above the left and right ports.

### Behavior

We trained rats to perform an auditory change detection task previously employed in studies involving human subjects (Booras et al., 2021; Ganupuru et al., 2019; Harun et al., 2020; Johnson et al., 2017). In this study’s implementation, rats inserted their nose into the central port of the behavioral apparatus cued by an LED light. Upon nose insertion, a stream of auditory pulses (“clicks”) was generated at a baseline rate of 20 Hz according to a Poisson process. Clicks were broadband and each had a duration of 3 ms. At a random point in time sampled from a truncated exponential distribution, with a mean of 2 s and a maximum time of 4 s, the generative click rate increased by a variable magnitude of 10 Hz, 23 Hz, 36 Hz, or 49 Hz, sampled evenly. The hazard rate for change times was flat. After a change occurred, the rat had 0.8 s to respond by withdrawing from the port. Successful withdrawal within the allotted time window was recorded as a “hit”, which was rewarded with a drop of water from a port on either the left or right of the central port. Failure to withdraw within the allotted time was recorded as a “miss” with reward withheld. Premature response in the absence of a change was recorded as a “false alarm” with reward withheld. Finally, on 30% of trials, no change occurred, in which case the rat was required to maintain fixation in the central port until the stimulus ended to achieve a “correct rejection”, which was rewarded. Sessions lasted for approximately 80 minutes. For behavioral criteria, we chose rats for surgical implants (see below) that had at least a 0.4 correct rejection rate and 0.5 hit rate on >300 trials per session.

### Electrophysiological recordings

Single-unit measurements were recorded with Neuropixel 1.0 probes (Jun et al., 2017; Steinmetz et al., 2018). Probe implants were assembled according to previous methods (Juavinett et al., 2019). A silver wire was soldered to the grounding contacts on the probe, and the probe was glued to a 3D-printed internal mount that was affixed to a stereotaxic adapter for implantation. The internal mount was then bound to an external mount by epoxy which additionally holds a headstage circuit board for data transmission.

Probes were stereotactically implanted between -3.80 mm and -3.95 mm AP relative to Bregma, +/- 2.2 mm ML relative to bregma (probe implanted contralateral to the side of the reward port assigned to the subject), and 1.5 mm ventral to brain surface (**Figure 2B**). Probes were lowered slowly, roughly 20 seconds per 0.1 mm, while simultaneously recording from them to ensure probe function and verify implantation depth. Ground wires were inserted directly into the cerebellum approximately 2 mm posterior to interaural zero (“IA0”). After implantation of probe and ground wire, both craniotomies were filled with sterile optical lubricant. Absolute Dentin was applied to the external mount of the probe and the ground wire to bind the implant to the skull, and dental acrylic was applied over the skull to seal the implant. The headstage was taped to the external mount with Kapton tape and the implant was covered with self-adhesive wrapping. Rats were left with free access to food and water to recover for one week following surgery before resumption of training and recordings.

During recording sessions, the self-adhesive wrapping was removed from the implant, and an interface cable was plugged into the headstage. The interface cable was wrapped around a gel toe sleeve to relieve strain from the rat moving (**Figure 2A**). The cable was fed through a simple pulley system to prevent the rat from grasping the cable while rearing, connecting to a PXIe acquisition module. This module interfaced with the Bpod and recording computer for temporal synchronization. Neural recordings were acquired using Open Ephys 3.

### Drug Infusions

Rats were implanted bilaterally with guide cannulas with a length of 4 mm at +/- 2.2 mm lateral and at either 3.80 or 3.95 mm posterior to bregma. Guide cannulas were lowered to the brain surface then cemented in place as described above for the probe implants. Dummy cannulas extending 0.5 mm past the tip of the guide cannula were inserted into the guide cannulas at the end of surgery. Following recovery from surgery and 3-4 days before the first infusion and data collection session, rats underwent a sham infusion in which they were lightly anesthetized with 2% isoflurane and internal cannula extending 1.5 mm past the tip of the guide cannula were inserted into the brain for four minutes. Rats were allowed to recover for 30 minutes before the behavioral session began.

Rats were infused twice a week before behavioral sessions with 0.3 µL of either saline solution (vehicle condition) or 1 mg/mL muscimol in saline solution (muscimol condition), alternatively. Volumes and concentrations were based on those of previous inactivation studies of this region, which were, in turn, based on autoradiographic and electrophysiological validation of muscimol spread dynamics (Martin, 1991). After being anesthetized with 2% isoflurane, rats were injected with either muscimol or vehicle through an internal cannula fitting inside the guide cannula and extending 1.5 mm past the tip of the guide. The internal cannula was filled with the saline or muscimol in saline solution and attached to a tube filled with mineral oil, which was attached on the opposite end to a Hamilton syringe used to control the injection. After slowly injecting saline or muscimol in saline solution over a period of two minutes, the internal cannula was left in the brain for four minutes to allow full diffusion of fluid. This process was repeated for each of the two cannulas. Like in the sham infusion, rats were left to recover for 30 minutes before the behavioral session began.

### Behavioral data analysis

False alarm trials included all trials in which rats withdrew their nose during stimulus baseline, including change and catch trials. Hit rates were calculated as the proportion of trials in which rats withdrew their nose within 0.8 s of the change point when a change actually occurred (i.e., excluding false alarm and catch trials). Reaction times for hit trials were calculated as the time of nose withdrawal after the change point. The influence of change magnitude on hit rates was quantified via logistic regression, and the influence of change magnitude on reaction times via linear regression with p-values derived from t-statistics applied to the fitted coefficients using the Matlab GLMFIT function. For analysis of the effects of muscimol on hit rates and reaction times, an additional regressor was included in GLMFIT as a shift term for the muscimol effect with p-values derived from t-statistics, similar to other regressors. Statistics on effects of muscimol on false alarm rates were determined using Fisher’s exact test.

Psychophysical reverse correlations (PRCs) were generated by aligning the click times of each false alarm trial to the time of the rat’s nose withdrawal. False alarm trials were used for PRCs to avoid the confound of changes in the generative click rate. For plotting the PRCs, each false alarm click time vector was convolved with a causal half-Gaussian filter with a standard deviation of 0.05 s and sampling every 0.01 s. Regardless of false alarm time and thus duration of stimulus preceding the detection report, all false alarms were included in the PRC, with false alarms with shorter stimulus durations simply contributing to a smaller epoch of the PRC. After this smoothing, false alarm reverse correlations were averaged together. Start points of false alarm PRCs were calculated using a fit of raw (i.e. non-smoothed) average click rates for each time point leading up to the PRC peak with a two-piece linear function with free parameters for base rate, start point of the upward slope, and slope. The Matlab FIT function was used to determine best fit parameter values and standard errors. The statistical comparisons of start points for the muscimol inactivation was performed with a t-test. Mean increases of the click rate were calculated over the interval from 50 to 350 ms before false alarms based on the raw click counts for each trial, only including trials that were at least 350 ms in duration. For PRC analyses comparing early and late false alarms, false alarm trials were categorized as “early” if the false alarm occurred within 1.5 s of stimulus start and “late” if the false alarm occurred later than 1.5 s after stimulus start.

### Electrophysiological data processing and analysis

We used the Kilosort 2.5 spike sorting algorithm (Pachitariu et al., 2016) to identify single unit clusters of spiking activity in our data to include for data processing.

Following initial sorting, the Phy 2.0 cluster viewing software was used to manually curate clusters to remove units with drop-out due to drift via visualization of amplitude plots over the course of a session. Phy also allowed us to merge clusters that clearly originated from the same unit by assessing correlation of spike times between clusters and correlations in drift. Units that drifted out of the probe’s recording range during a session were excluded from analysis to avoid distortion of trial-by-trial response observations.

We further filtered single units for inclusion in analysis by identifying units with at least a 1 Hz firing rate during active task engagement (i.e. during stimulus presentation and reward collection). The remaining units were again filtered for task modulation by comparing firing rates on hit trials between stimulus start and change time to the firing rates between change time to the time of response (two-sample t-test, p < 0.05). Units were sorted into two groups: those with a higher firing rate during baseline, comprising negatively modulated units, and those with a higher firing rate following stimulus change, comprising positively modulated units.

Peri-stimulus time histograms (PSTHs) were calculated by aligning spikes to one of two stimulus events: time of the change or decision report nose withdrawal (“response”). In both cases, spikes were convolved with a Gaussian filter (0.1 s standard deviation). For each unit, neural activity was then z-scored based on the trial-by-trial firing rates on hit trials between stimulus start and change time. Neural traces were sorted based on the programmed change magnitude (+10, +23, +36, +49 Hz). For traces aligned to the change, plots were extended to the median reaction time for each condition and included no part of any trial after the decision report nose withdrawal. For traces aligned to the decision response, only hit trials were included.

Linear regression was used to test for modulation by change magnitude based on a linear model of z-scored firing rate with a shared offset for all conditions and a single slope regressor for change magnitude. For neural activity aligned to the change, the regression was performed on the z-scored firing rates in the interval from 100 to 200 ms after the change. For neural activity aligned to the decision response, the regression was performed on the z-scored firing rates in the interval from 150 to 50 ms before the decision response. The Matlab function REGRESS was used to perform this analysis and determine standard error for the regressor coefficient and p-values based on a t-statistic.

Click-triggered averages (CTAs) were calculated by aligning neural responses to the time of each click. To avoid non-causal influences of clicks on neural activity, spikes were convolved with a causal half-Gaussian filter (0.1 s standard deviation) for the CTA analyses. As in the PSTHs, neural activity was z-scored for each unit based on the trial-by-trial firing rates on hit trials between stimulus start and change time. Then, for each trial, the average activity for the corresponding unit was subtracted to obtain the “residual neural activity” on a trial-by-trial basis – that is, how the neural activity at any point on a trial differed from the average activity at the same time point for the corresponding unit. This removed the average time-dependent modulation, including contributions expected from other clicks. The residual neural activity was then aligned to the time of the onset of each individual click. This analysis was focused solely on clicks that occurred in the baseline period before any change in the generative click rate to avoid any confounding influence of that change. For comparing CTAs occurring early or late in trials, we included responses to clicks occurring earlier than 1 s from stimulus onset in the “early” category and responses to clicks occurring later than 1 s from stimulus onset in the “late” category. This separation was chosen to be 500 ms earlier than that used for the PRC analysis to account for the fact that the CTA shows the influence of a click forward in time while the PRC shows the influence on choices of clicks that occurred earlier in time, with a multiple hundred millisecond “integration time” implied by both. Comparison of CTAs for early versus late clicks for individual neurons (**Figure 4C**) were based on the maximum amplitude of each neuron’s CTA in the first 500 ms after click onset. CTAs for negative-modulated units focused solely on later clicks that occurred at least 1 s after stimulus onset, corresponding to the clicks that had the most robust influence on the positive-modulated units. Statistics for all cases were calculated based on the CTA magnitudes in the interval from 50 to 350 ms after click onset with p-values based on a t-statistic.

## Acknowledgements

We thank Christine Angeles and Ani Tunnell-Braun for assistance with animal training and Aya Akhmetzhanova for helpful discussions. PG, TS, and KS were all supported by the Training Program in Basic Neuroscience T32 (MH082174) and the Learning, Memory, and Plasticity (LAMP) T32 (MH112507). TS was additionally supported by an ARCS Foundation Scholar Award. KS was additionally supported by the NeuralStorm NSF NRT training grant (NSF 2152260). RC and TDH were supported by a collaborative CRCNS grant (NSF 2207895) and a UC Davis Memory and Plasticity Seed Grant. TDH was additionally supported by a National Alliance for Research on Schizophrenia and Depression Young Investigator Award, a Whitehall Foundation Grant, and the National Institutes of Health (NIH R21 MH114147, NIH R56 MH120283, NIH R01 MH124818).

